# Assessment of Total Oocyte Transcripts Representation through Single Ooplasm Biopsy in Bovine with High Reliability

**DOI:** 10.1101/2023.04.24.538116

**Authors:** Dewison Ricardo Ambrizi, Ricardo Perecin Nociti, Tiago Henrique Camara De Bem, Joao Vitor Puttini Paixao, Jacinthe Therrien, Elisangela Chicaroni De Matos, Jose Bento Sterman Ferraz, Marcos Roberto Chiaratti, Juliano Sangalli, Juliano Coelho Da Silveira, Felipe Perecin, Lawrence Charles Smith, Flavio Vieira Meirelles

## Abstract

Understanding the entire transcriptional and epigenetic landscape is facilitated by the application of omics in a number of ways. Today, omic instruments are more affordable and easier to implement. In human research, for instance, single-omics are a reality and are used extensively to generate vast quantities of data. This method permits the comprehensive reconstruction of transcriptome and epigenetic markers removing bias from pooled samples. In tandem with the evolution of machines and protocols, algorithms and genome annotation have undergone continuous improvement. The genome annotation of domestic animals is inferior to that of humans, rodents, and less complex organisms. In the case of heifers, the reference is incomplete, with significant gaps and only a portion of the noncoding transcripts. The purpose of this study is to validate our compartmentalized single oocyte biopsy by comparing a small fraction of bovine oocytes, 1%, to the entire oocyte at the Metaphase II stage. In addition, we examined the use of four database sources (NCBI, ENSEMBL, UCSC, and NONCODE) to produce a merged non-redundant gene alignment and counting in order to enhance gene detection and normalization, resulting in a more accurate method to comprehend the entire landscape. This study is a continuation of our research titled “**Retrospective model utilizing biopsies, granulosa cells, and polar body to predict oocyte competence in bovine**” in which this method was used to retrospectively compare biopsy oocytes collected during the MII phase. With the addition of NONCODE information, gene normalization was significantly enhanced. In addition, our analysis identified 4560 noncoding genes from NONCODE references. ENSEMBL and NCBI have nearly the same number of annotated genes (16,423 vs. 17,804), but using ENSEMBL as a reference, 2356 genes were able to be normalized and identified. Proceeding to biopsy x oocyte analysis, we were able to detect a greater number of genes in oocytes than in biopsy, where the preponderance was from NONCODE sources (68). Despite these minor differences, the high correlation of expression between them (89%) was consistent and proved to be a valuable instrument for studying the oocyte without destroying it.

## INTRODUCTION

The omic atlas from several tissues, species, and conditions has been continuously expanding over the past few years through technological improvements, generating data with more coverage than ever. Additionally, these omic tools are constantly becoming cheaper, leading to a logarithm increase in the amount of generated data (Chen et al., 2021; Lyfer et al., 2023). The widespread use of new technology is a normal process that all technologies undergo, beginning with high costs and complexity and gradually becoming cheaper and more accessible for large-scale implementation. Technological advancements, particularly in single cells, have made omics data a suitable tool to understand the entire cellular landscape, cell by cell, or from small and limited samples using different techniques to achieve specific goals.

Nowadays, the single-cell approach (Sc) can be used for a variety of purposes with different omic techniques, such as investigating differentially expressed genes with less bias and tissue specificity (RNA-Seq), protein-chromatin interaction (Chip-Seq and CUT&RUN), chromatin accessibility (ATAC-Seq), DNA-methylation (methyl-Seq or whole genome bisulfite sequencing), chromosome conformation (Hi-C), and finally, use all these generated data in an integration analysis (Islan et al., 2011; Saliba et al., 2014; Telenius et al., 2018; Hainer et al., 2019; Cusanovich et al., 2015; Guo et al., 2013; Nagano et al., 2013).

Recently, Sc-RNA-seq has become one of the most popular Sc-omics. It includes the study of cell and subcellular populations in a tissue-specific manner, cancer populations, and sequencing of different embryo layers and subcellular types in the domestic animal reproduction field. While pooled RNA-seq is still prevalent in animal sciences, Sc-RNA-Seq is ascending, especially in reproduction. The advantages of these Sc approaches are the efficiency and accuracy of these new technologies and, most importantly, the opportunity to design complex experiments to answer old questions that were not previously possible due to technical limitations. Moreover, single omics can avoid pooled samples’ bias and add the natural and genuine biological variations intrinsic to data, which is indispensable for extracting the true information and reaching the real interpretation.

Concomitant with improving the sequencer equipment, the algorithms tools for alignment, trimming, quality checking, and statistical models to find significance in data also have advanced. Together, genome annotation has been constantly updated in domestic animals, increasing the accuracy, the amount, and the classification of transcripts. This improvement impacts each transcript class’s function and biological process, as evidenced by the FAANG consortium (Giuffra et al., 2019). This consortium emphasizes the importance of genome annotation variability in omic analysis and annotation reference, as the primary bias in the old annotation in bovine was the absence of variation. Another example of a genome annotation source is the NONCODE consortium, which has increased the number of transcripts that in the past were neglected (Bu & Yu, 2011).

In this study, we are motivated by three different contexts:

1. We aim to estimate the correlation between the transcripts present in the MII oocyte biopsy and the respective donor oocyte.
2. We plan to amplify the annotation source using ENSEMBL and NCBI removing the redundancy between them and still, adding the NONCODE information for the oocyte transcriptome.
3. We intend to study the maternal RNAs in the ooplasm (MII) and correlate them with the competency of development, avoiding destroying the oocyte.

To verify the feasibility of the third context, we conducted an experiment comparing bovine oocyte (MII) biopsies with the respective donor oocyte using the Sc-RNA-Seq approach for each sample as described in Figure 1A. We included low-counting genes and exclusive genes, usually left behind in standard analysis using a more accurate and complete way to align and count genes. As result, we found that transcriptome profiles comparing biopsies and oocytes are highly correlated. The variance between conditional samples (oocyte or biopsy) was higher than the present in paired samples. Moreover, we detected sixty-eight genes exclusive from the oocyte and not in the biopsy. Additionally, we identified many non-coding transcripts, suggesting that these RNAs perform an essential regulatory role at this stage of development. These results indicate that the method performed in this work is feasible and can be used to compare oocyte and biopsy with high reliability, since it was able to detect more RNAs (genes, non-coding, etc.), improving the robustness and quality of the analysis.

**Figure 1.**
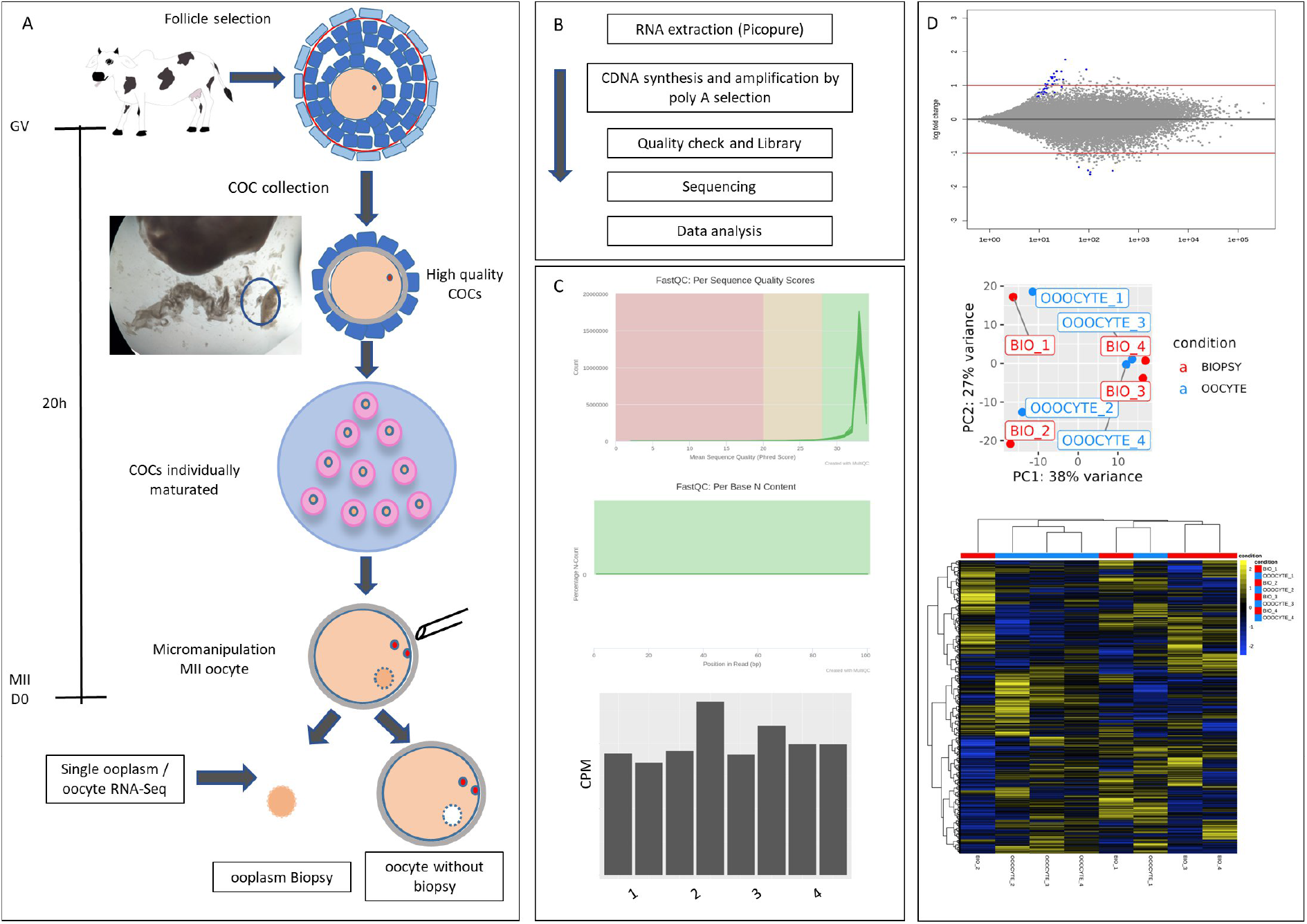
Experimental design and sequencing. A - Follicles with five-millimeters were manually dissected from ovaries collected from an abattoir. The COCs were selected and *in vitro* maturation was performed for 20h. After identifying MII oocytes with 1° PB extrusion, an ooplasm biopsy was performed and both the biopsy and biopsied oocyte were snapped and frozen in separate tubes. A total of four oocytes and the respective biopsy were collected. B - Total RNA was extracted using Picopure and then immediately converted to cDNA. The quality was checked using Bioanalyzer and then prepared in the library. Sequencing was performed with 10 × 10^6^ reads per sample. C - Phred Score and Per base N Content quality. A trimming process was performed to remove base calls with insufficient confidence. The counts per million for each sample pair (biopsy x oocyte) were stable among the samples. D - DESeq2 log2FoldChange of biopsy x oocyte comparison from the mean of normalized counts for all samples. The PCA reduction dimension method showed an accumulation of 38% in PC1; heatmap using ward.D2 was performed as another clusterization method.

## RESULTS

### Sample sequencing

Our first question was related to RNA sequencing quality and whether the amount of biological material in the ooplasm biopsies was enough to obtain high-quality sequences. Briefly, we estimated the ∼volume of the oocyte biopsy once we know the oocyte volume and the internal diameter of the injection needle. Considering that the oocyte has ∼110µM of diameter, *4/3* x *Π x r*^*3*^ resulted in 696.556,666um^3^. In addition, the internal diameter of the injection needle was calculated assuming that it was a perfect cylinder, *Π* x *r*^*2*^ *x h* resulting in 10602.875 um^3^ (*Π x 7*.*5*^*2*^ *x 60µm*). This amount is a ratio of 0.0150 (1.5% of ooplasm), being fairly superior to a single cell volume of oocyte biopsy, which suggested to us the feasibility of the technique. After sample processing and amplification, we performed a quality check step and then samples with RIN > 0.7 were sequenced using Illumina plataform (Figure 1B). Sequencing quality was performed and the Initial steps to examine the quality of the sequencing reads provided the Phred Score and Per base N Content (Figure 1C), indicating more than 99% of precision (Phread score > 30) and high quality after trimming for GC percentage and length (Table 1).

**Table 1.**
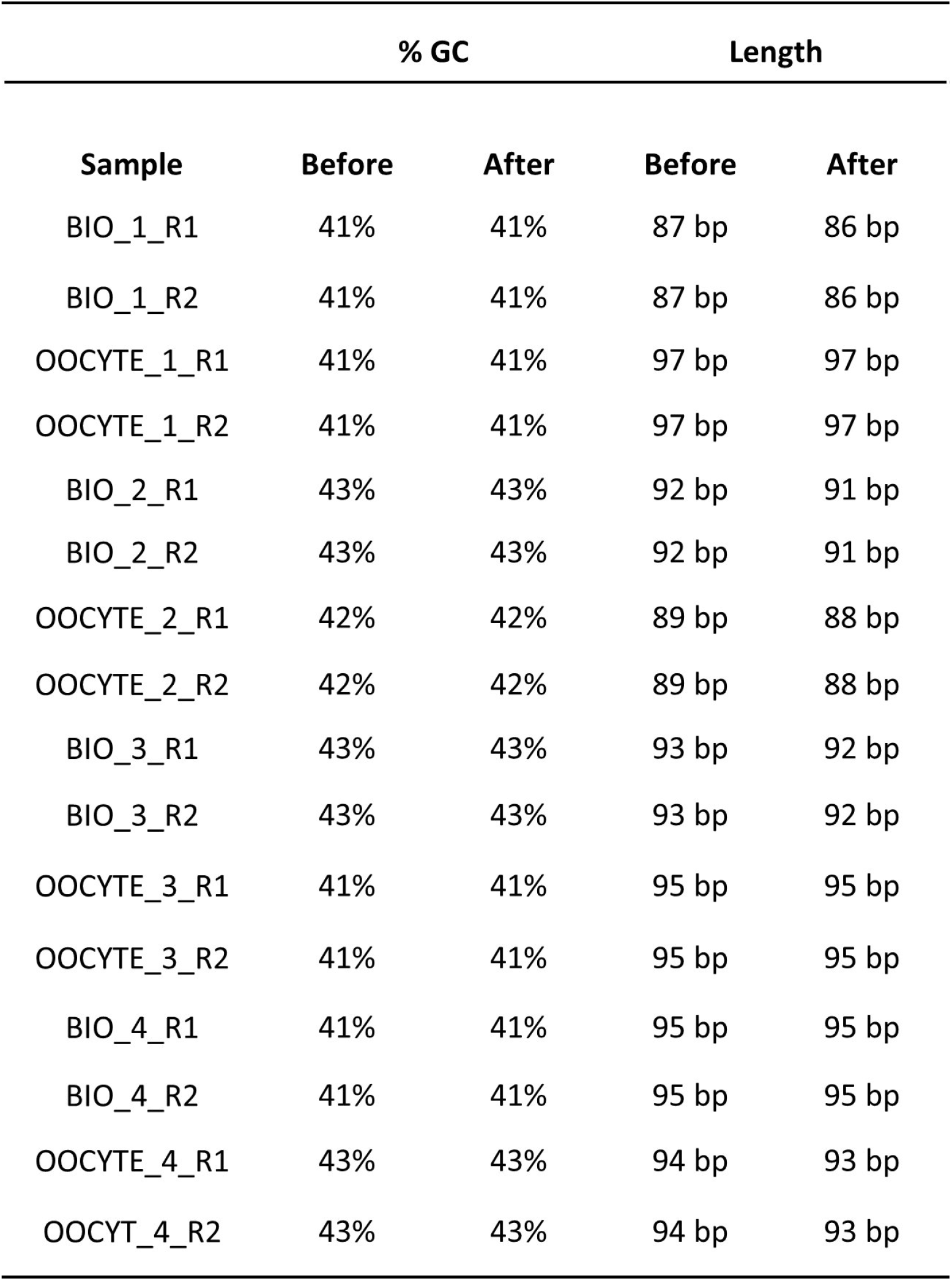
Quality check comparing before and after trimming process.

Despite the removal of 20bp, there was no significant decrease in bp length and the percentage of alignment reads was higher when compared with the alignment using untrimmed data, probably a consequence of the removal of short reads (less than 30bp). The counts per million (CPM) for each sample pair (oocyte X biopsy) were stable among the samples, indicating that our approach efficiently amplifies and normalizes the total amount of transcripts sequenced. For gene normalization, we only considered genes with a minimum of three counts in at least one of the groups.

As indicated in the MA plot (Figure 1D), the mean of normalized counts in oocyte x biopsy comparisons showed that only a few genes deviated from the log2Foldchange interval and were positioned outside the confidence interval (adjusted P higher than 0.1). Total variance, as indicated by the principal component of variance analysis (PCA), indicates that the PC1 accumulated 38% of the variance. Interestingly, the oocyte samples clusterized as pairs with their respective biopsy pairs and not according to condition (Figure 1D). This result strongly indicates higher similarity between pairs than among the sample condition (oocyte or biopsy), suggesting the variation is associated with differences between oocytes. Finally, we performed a heatmap clusterization analysis (Figure 1D), another reduction dimension method to check how the VST variance Stabilized data could be arranged. This graphical visualization identified a similar result found in the PCA, where the majority of samples were not grouped by condition.

### Improving bovine gene detection and normalization

RNA-Seq is a highly precise and informative method for analyzing gene expression, with a hundred-fold greater accuracy compared to qPCR. However, several factors can affect RNA-Seq, such as read quality, fragment size, alignment tool, and the number of genes. For instance, the DESEq2 method performs internal normalization by calculating the geometric mean for each gene across all samples. The counts for a gene in each sample are then divided by this mean, and the median of these ratios in a sample is the size factor for that sample. Therefore, the number of genes directly influences the normalization process, which can impact the discovery of real differences in gene expression.

Moreover, annotation sources, such as ENSEMBL, NCBI, and UCSC, provide a great number of genes and transcripts, which sometimes do not overlap, adding bias according to the genome reference used. To address this issue, we used the ENSEMBL, NCBI, and UCSC together with NONCODE databases to improve our knowledge of annotated non-coding genes. We used the STAR algorithm as recommended by nf-core protocols.

To determine how much information could be added to the annotation process established by comparing the three databases, we conducted a general overview and aligned the databases using ENSEMBL as the base for removing redundancy. Our analysis revealed that some RNA biotypes were only found in ENSEMBL, including immunoglobulin, mitochondrial rRNA and tRNA, processed pseudogene, pseudogene, ribozyme scaRNA, sRNA, and T-cell receptor (TRs) (Figure 2, Table 2). Conversely, antisense-RNA, tRNA, C-region, and V-segment were only present in NCBI due to their exclusivity to this database. Additionally, we found that misc-RNA and protein-coding biotypes were present in both databases proportionally, while miRNA, rRNA, snoRNA, and snRNA had more genes in ENSEMBL than from NCBI.

**Figure 2.**
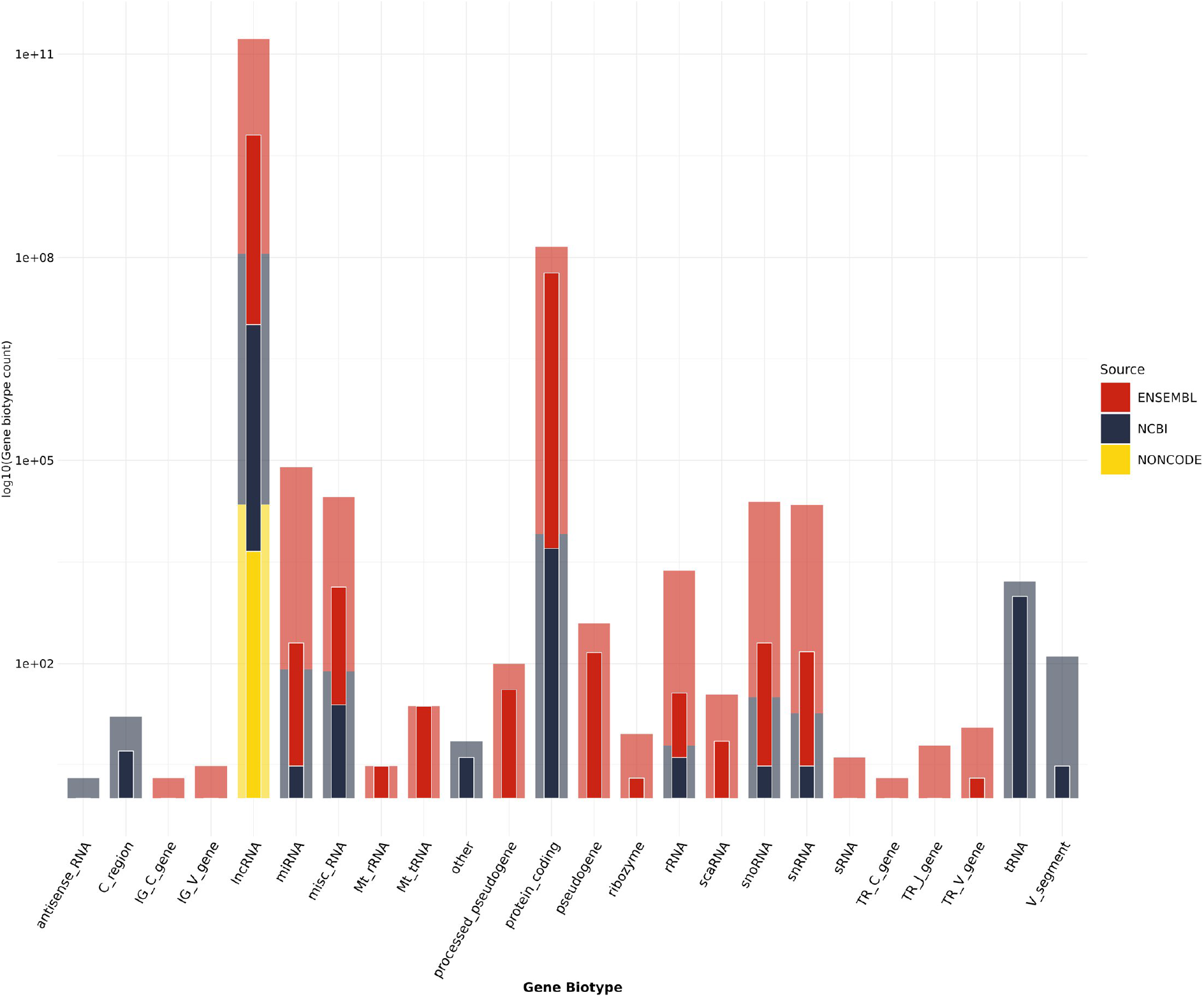
Counting gene biotypes identified in oocyte and biopsy sequencing using the SC-RNA-Seq approach. The Y axis is represented in a log10 scale. Shaded bars represent the total RNA biotype annotated, while dense bars (front) show the amount found in our data. Colors represent different repository sources.

**Table 2.** Tabular values clustered by RNA class and annotation source.

| gene_biotype | Source | Annotated | Identified |
| --- | --- | --- | --- |
| C_region | NCBI | 15 | 4 |
| IG_C_gene | ENSEMBL | 1 | 0 |
| IG_V_gene | ENSEMBL | 2 | 0 |
| Mt_rRNA | ENSEMBL | 2 | 2 |
| Mt_tRNA | ENSEMBL | 22 | 22 |
| TR_C_gene | ENSEMBL | 1 | 0 |
| TR_J_gene | ENSEMBL | 5 | 0 |
| TR_V_gene | ENSEMBL | 10 | 1 |
| V_segment | NCBI | 127 | 2 |
| antisense_RNA | NCBI | 1 | 0 |
| lncRNA | ENSEMBL | 1,470 | 629 |
| lncRNA | NCBI | 5,102 | 2,235 |
| lncRNA | NONCODE | 22,227 | 4,560 |
| miRNA | ENSEMBL | 965 | 66 |
| miRNA | NCBI | 81 | 2 |
| misc_RNA | ENSEMBL | 372 | 55 |
| misc_RNA | NCBI | 75 | 22 |
| other | NCBI | 1 | 1 |
| processed_pseudogene | ENSEMBL | 99 | 40 |
| protein_coding | ENSEMBL | 17,579 | 11,788 |
| protein_coding | NCBI | 8,111 | 4,975 |
| pseudogene | ENSEMBL | 389 | 145 |
| rRNA | ENSEMBL | 393 | 8 |
| rRNA | NCBI | 5 | 3 |
| ribozyme | ENSEMBL | 8 | 1 |
| sRNA | ENSEMBL | 3 | 0 |
| scaRNA | ENSEMBL | 33 | 6 |
| snRNA | ENSEMBL | 1,217 | 49 |
| snRNA | NCBI | 17 | 2 |
| snoRNA | ENSEMBL | 787 | 66 |
| snoRNA | NCBI | 30 | 2 |
| tRNA | NCBI | 57 | 12 |

Our analysis showed that the NCBI source provided an additional source of information to improve annotation, contributing 9170 genes to our annotated gene analysis. Furthermore, the NONCODE database contributed an additional 22,227 genes. Among the annotated genes, we were able to add 8286 genes (3717 from NCBI and 4569 from NONCODE) that were previously undetected when only ENSEMBL was used. This demonstrates the importance of incorporating multiple annotation sources to improve the accuracy and completeness of the bovine genome annotation. Proportionally, oocytes accumulate 27.7% of all annotated genes and 37.08% compared to ENSEMBL and NCBI, the most popular databases. The numeric values can be visualized in Table 2.

We also compared the individual effect of ENSEMBL or NCBI in gene normalization and in the discovery of differences between groups. We found a large overlap between ENSEMBL and NCBI in terms of genes found and normalization, demonstrating that the total number of genes is almost the same (16,423 vs. 17,804), with some genes present only in one annotation and vice versa (Figure 3A). However, a higher density was observed for ENSEMBL in the middle of the distribution, suggesting a difference in transcripts with high counts in ENSEMBLE that were not found in NCBI, despite its higher number of total genes. The merged approach applied here enabled a higher gene detection, increasing the total genes with a smaller basemean when it is compared with ENSEMBL and NCBI databases. Part of this skew is due to the low-count genes that were added with the NONCODE source. However, 4560 genes were added from NONCODE and the difference of the merged approach with ENSEMBL and NCBI was 8297 and 6916 respectively, indicating that the addition of lncRNAs in the normalization process added 3737 and 2356 genes, respectively.

**Figure 3.**
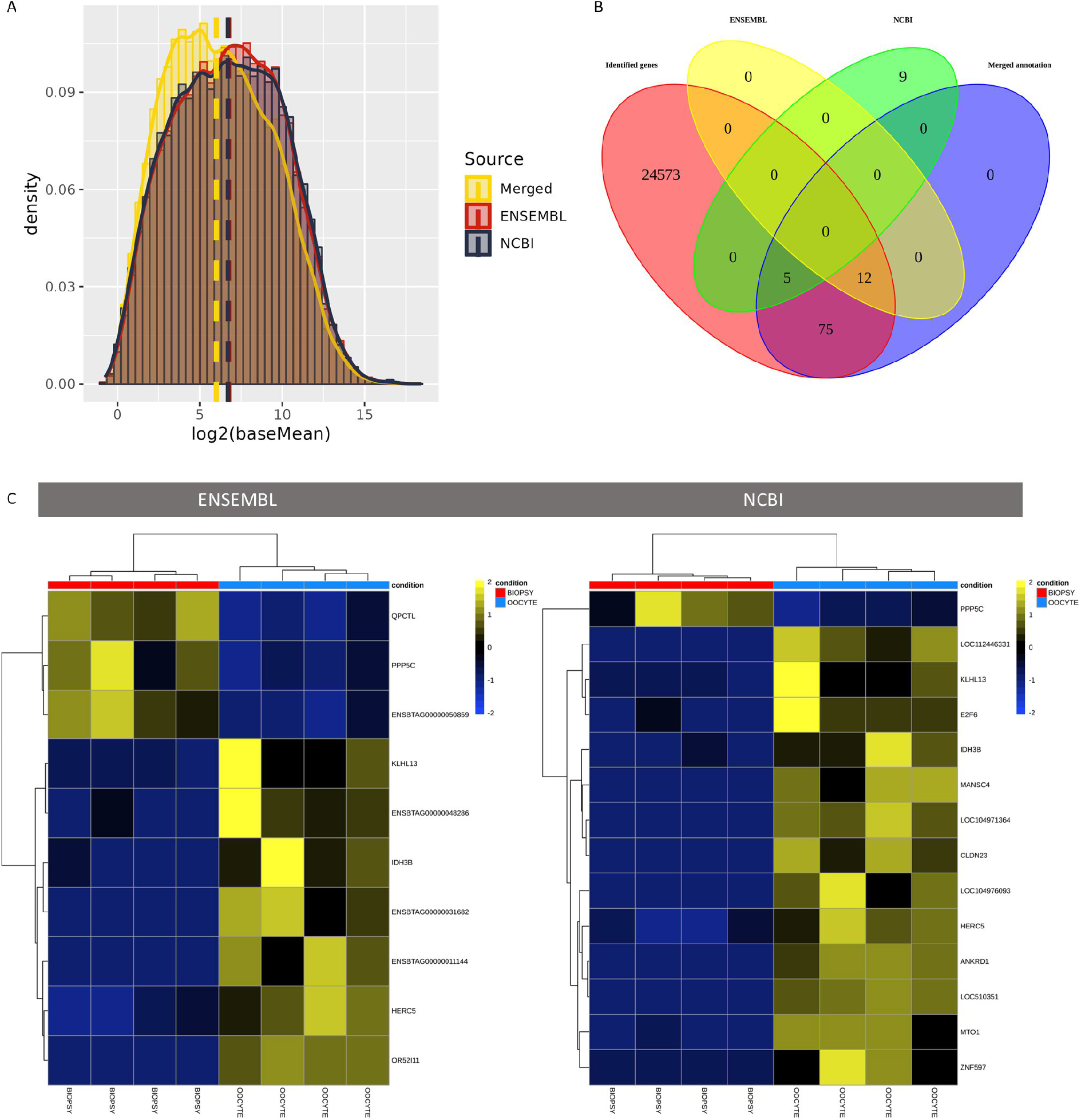
Differences using only one annotation or combined annotation source. A - Log2(basemean) showing the differences in distribution comparing only ENSEMBL, NCBI or merged annotation. B - Venn diagram showing differences in gene expression. C - Heatmaps for differences in Biopsy x Oocyte comparison based on the ENSEMBL or NCBI annotation, respectively.

Differences in normalization, even in data with low variation between groups such as ours, could detect different genes in comparison (Figure 3B, C). Using the nomenclature of the TPM table, we identified some significant differences between oocyte and biopsy. The ENSEMBL source had more genes in common with merged annotation in comparison with NCBI (12 × 5). In addition, 75 genes were only found when the merged annotation was used, suggesting a better normalization since the added genes brought the data closer to the real transcriptome landscape.

Proceeding with the merged annotation, we identified more differences and more genes besides the lncRNAs added from the NONCODE source. This approach enabled the discovery of more DEGs and exclusive genes (more than one count per sample in at least three samples from one group) (Figure 4A) once it had an impact in gene normalization. Interestingly, between the DEGs found in the oocyte and biopsy, we identified a higher variability in those upregulated in the oocyte than those upregulated in the biopsy (Figure 4B), suggesting a higher variability in the oocyte than biopsy for DEGs genes. However, the gini index for oocyte-exclusive genes was low for most genes, indicating a lower variability in counts. DEGs and exclusive genes are displayed in heatmaps (Figure 4C). In general, more lncRNAs from NONCODE sources and other genes are likely to be localized around the oocyte’s MII spindle and would be absent in ooplasm biopsies obtained far from the spindle, suggesting the intracellular compartmentalization (heterogeneity) of some transcripts within the oocyte.

**Figure 4.**
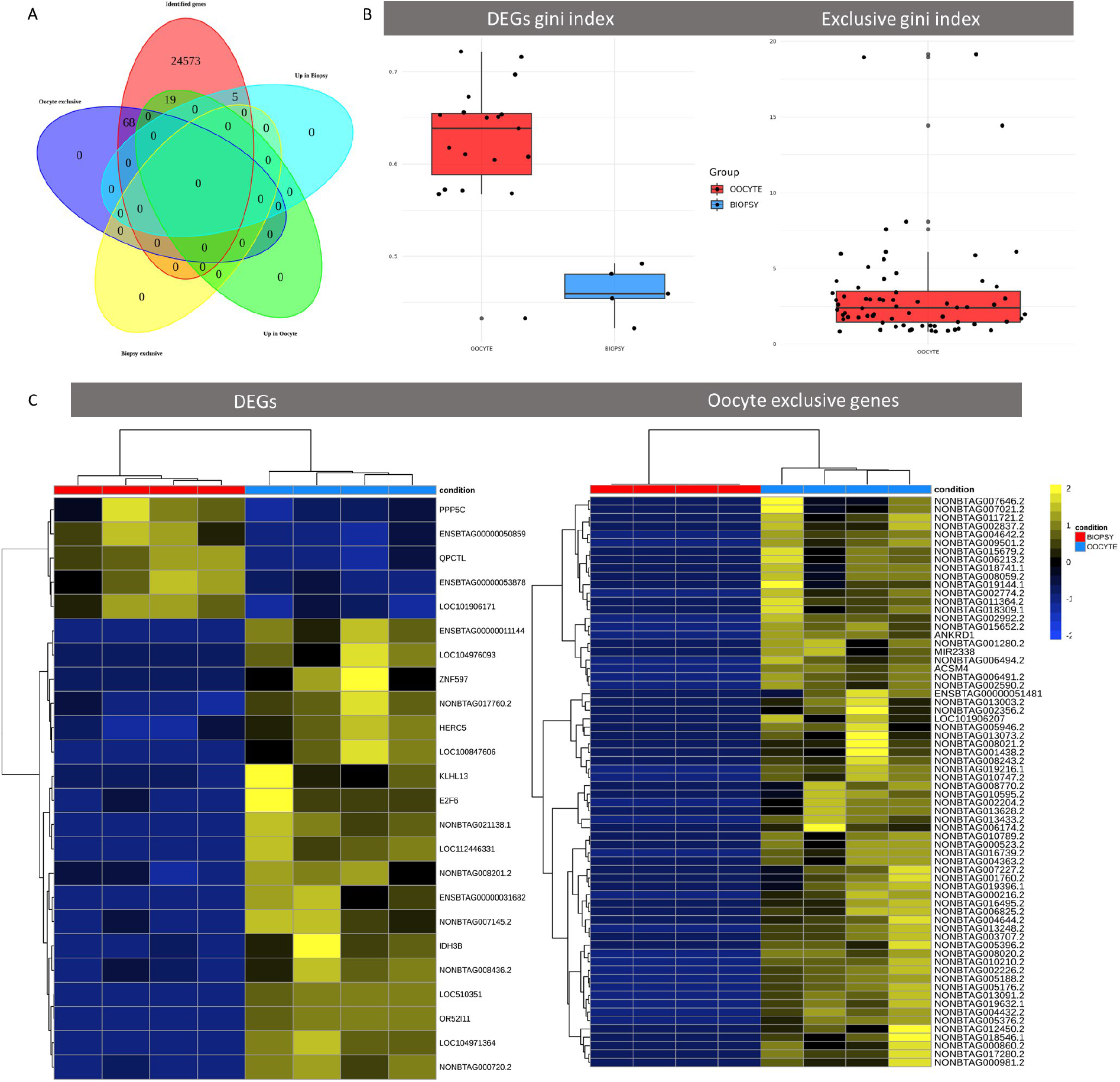
Exclusive genes analysis. A - Venn diagram showing differences found between biopsy and oocyte. In a total of 24646, 68 were found only in oocytes as exclusives; 19 and 5 were found as DEGs respectively for oocyte and biopsy. B - Gini index for exclusive and DEGs genes found in biopsy and oocyte comparison. C - Heatmaps of DEGs and exclusive genes in oocytes.

Some studies involving mammalian oocyte RNA localization indicated that there is heterogeneity in the GV phase, but not in MII (Jansova et al., 2021). Still, the heterogeneity found in the GV phase is linked to a recently discovered membrane-less structure named MARDO (mitochondria-associated ribonucleoprotein domain) which also includes RNAs such as ZAR1 and YBZX2 before the MII phase. However, this MARDO is dissociated during the transition from MI to the MII phase (Cheng et al., 2022). We did not find differences in these genes in the biopsy x oocyte comparison, which was an expected result once only oocytes with the first polar body were used in this experiment.

## CORRELATION ANALYSIS

Correlation co-expression analysis in RNA-Seq is a computational method used to investigate the relationship between gene expression levels across samples. Gene expression is typically quantified as the number of reads that map to each gene in a given sample. By comparing gene expression levels across samples, we can identify genes that are co-expressed, meaning that their expression levels tend to increase or decrease in a similar manner.

Here, we performed the log2 correlation using the rlog function, similar to variance Stabilizing Transformation (VST) from the DESeq2 package. Results indicated a high correlation (> 0.98) among paired oocyte-biopsy samples (Figure 5A). In addition, the Spearman correlation found similar results with a minimum and maximum of around 0.86 and 0.91 (Figure 5B). This difference in results can be attributed to the normalization method of the rlog function, which corrects better for the small values in the normalization process. Moreover, for Spearman correlation analysis, the RNA-Seq of granulosa cells from our database was added to show that the correlation found between oocyte and biopsy was not random. Correlation among granulosa samples was around 0.82-0.96, but comparing granulosa with oocyte or biopsy samples, we found a correlation interval of 0.4-0.59 (Figure 5B). This is an indicator of co-expression correlation based on the similarity of gene expression that suggests a non-random correlation value found in our samples (oocyte vs. biopsy). Reinforcing this result, using the log2 TPM for each group, the dispersion between granulosa x biopsy, and granulosa x oocyte was graphically performed (Figure 5C). These visualizations evidently showed that the dispersion between granulosa and biopsy or oocyte was extremely variable, while oocyte x biopsy dispersion was an almost perfect match with all the points from both groups under the line showing a perfect linear distribution (Figure 5C).

**Figure 5.**
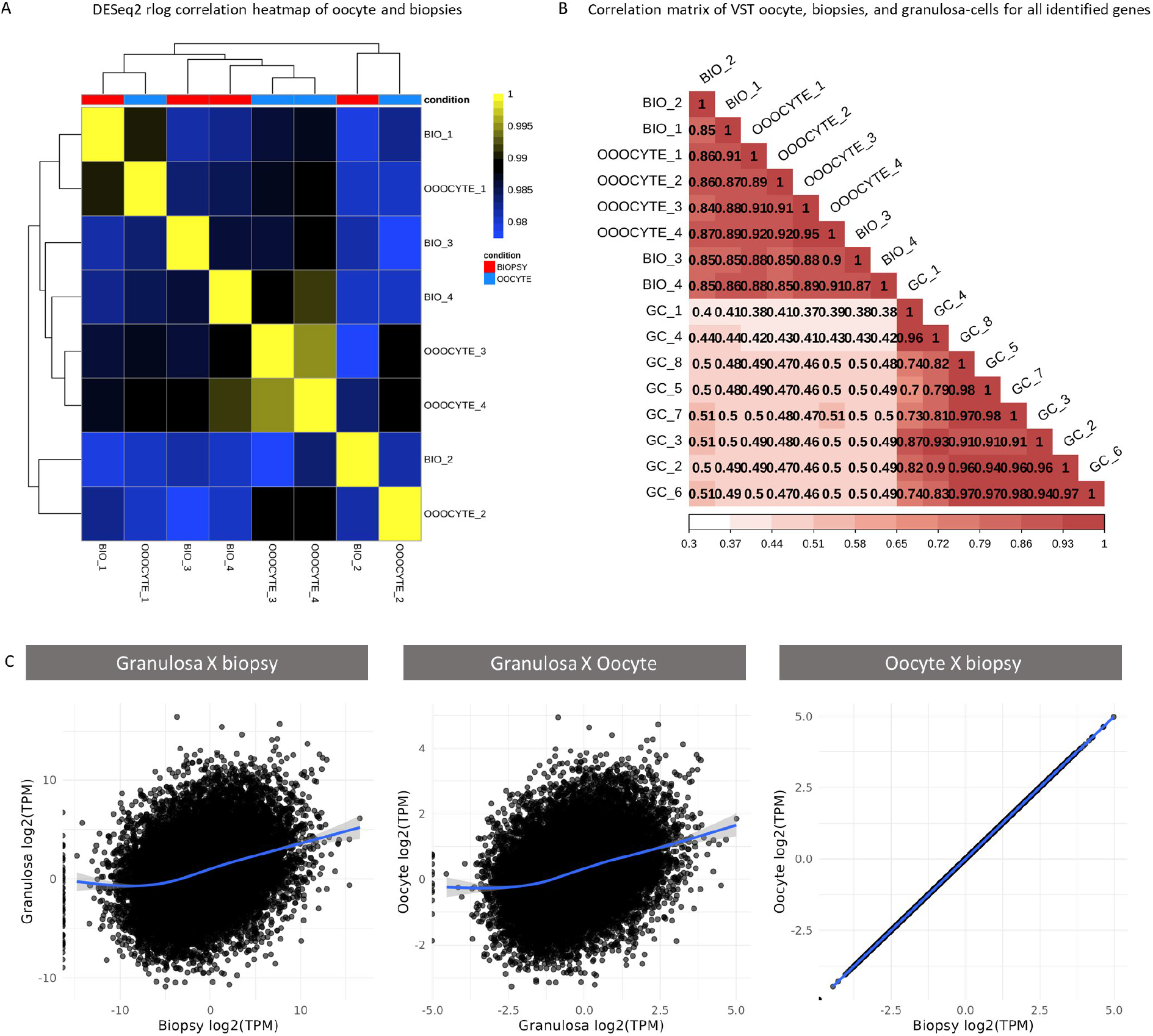
Correlation analysis. A - Heatmap correlation was performed using the DESEq2 normalized object using the rlog function. B - Correlation matrix comparing oocyte, biopsy and granulosa cells (negative control) data sequenced at the same conditions of biopsy and oocytes. C - Dispersion point graphic using Log2 TPM values for granulosa x biopsy, oocyte x granulosa, and oocyte x biopsies showing the relationship between each comparison.

## CONCLUSION

Mammalian oocytes store a large quantity and variety of RNAs in the ooplasm, which gradually accumulates during oocyte growth and is directly correlated with the growth of follicles. These maternal RNAs are essential to development, mainly during the oocyte-zygote transition. However, they are consumed (degraded or translated) during embryo development, making it difficult to relate the function of some RNAs with oocyte competence.

The oocyte volume (um^3^) is approximately 2000 times larger than a fibroblast and 133,333.33 times larger than a sperm. On the other hand, an ooplasm biopsy contains ∼3500um^3^ (based on the tubular volume area of the injection pipet), representing ∼ 2.7% of the ooplasm volume, which is still fifty times larger than a cell. Therefore, the methodology is feasible even considering technical limitations.

Summarizing, our results indicate that the method performed in our work was feasible to compare the content of RNAs in biopsies and oocytes with high reliability, improving the robustness and quality of the analysis. In a general manner, we found some small differences between biopsies and oocytes. However, the results (PCA clusterization) showed that each biopsy was clustered with its respective oocyte, indicating that the variability from the samples could be attributed to oocyte variation and not biopsy vs. oocyte variation. Still, this suggests that oocytes from high-quality COCs can have a distinct transcriptional profile, which is in line with our knowledge about oocyte growth and development, given that only a small fraction of oocytes have the true potential to develop until the blastocyst stage. Moreover, the differences found in DEGs presented a high Gini index variation, suggesting variability of these genes, which can be noisy and not a true variation. Furthermore, we would not have found these genes if we performed a normal analysis using only ENSEMBL or NCBI annotations. The majority of exclusive genes came from the NONCODE database, which, besides adding more genes from its own source, interfered positively with gene normalization, improving the discovery of other genes in ENSEMBL and NCBI sources. Finally, our data presented here suggest that ScRNA-Seq from oocyte biopsy is a valuable representation of the RNAs present in the oocyte due to its high fidelity and accuracy.

## MATERIAL AND METHODS

### Oocyte collection and sample preparation

Ovaries were collected from a local slaughterhouse and transported to the laboratory in saline solution (0.9%) with antibiotics (penicillin and streptomycin x100) at 37ºC. The follicles were dissected with the aid of sterile scissors and scalpels and the size of the follicles was measured with an accurate pachymeter. Only follicles with 5mm were used for the experiment. The dissected follicles were maintained in a physiologic solution with antibiotics. After, the follicles were ruptured, one by one, with a needle (40×12) assistance under a stereomicroscope. The immature oocytes were collected and washed three times in maturation media and posteriorly the *in vitro* maturation was performed. The oocytes were matured in micro-droplets (10μL), with one oocyte per droplet (single oocyte). We adopt this protocol to make this experiment compared to the retrospective experiment (**Retrospective model utilizing biopsies, granulosa cells, and polar body to predict oocyte competence in bovine**) previously published for our group. The maturation was performed in TCM-199 medium (ThermoFischer) supplemented with 10% of fetal calf serum (FCS), 5.0 μg/ml luteinizing hormone (Lutropin-v, Vetrepharm), 0.5 μg/ml follicle-stimulating hormone (Folltropin-v, Vetrepharm), 0.2 mM pyruvate and 50 μg/ml gentamicin under mineral oil at 38.5°C and an atmosphere of 5% CO2 in the air. After 19h of maturation, the 1^st^ polar body was selected, for this the cumulus cells were removed with a stripper pipet in an enzymatic solution (TripleExpress®). In addition, we used only MII oocytes with good morphological and cytoplasm characteristics.

### Ooplasm Biopsy

The MII oocytes were washed three times in TCM 199 Hepes supplemented with 10% of FCS, 0.2 mM pyruvate and 50 μg/mL gentamicin and kept, individually, per 15 min in micro-droplets of TCM Hepes with 7.5 μg/mL cytochalasin B. Microsurgery was performed using borosilicate micropipettes (holding and injection pipettes – Eppendorf, Germany) coupled at an inverted microscope (Nikon Eclipse Ti), equipped with micromanipulators and microinjectors (Narishige, Tokyo, Japan). Polar body was used as a reference point aligned at the 3 hours position. Then, all biopsies were removed from the same area at 5h position avoiding metaphase remotion. All biopsies were removed with the same ooplasm volume. Following the microsurgical recovery of the ooplasm biopsies and the corresponding biopsied oocyte were washed in PBS containing RNAse-out (ThermoFisher) as recomended to improve RNA quality and placed individually in separate low binding tubes. Samples were immediately snap frozen in liquid nitrogen (LN_2_) and storage at -80ºC for further analysis.

### RNA amplification

cDNA synthesis and amplification were performed using the SMART-Seq HT Kit (Takara Bio Inc), following the manufacturer’s recommendation. Next, cDNA was purified using AMPure XP Beads (Beckman Coulter), and cDNA concentration and profile were determined using Qubit dsDNA High Sensitivity (ThermoFisher Scientific) and Bioanalyzer High Sensitivity DNA Kit (Agilent), respectively. Libraries were prepared using the Nextera XT DNA Library Prep (Illumina) following the recommendations from the SMART-Seq HT kit. Libraries were assessed using Qubit and Bioanalyzer, before sequencing on the NextSeq 2000 (Illumina) considering 100 bp paired-end reads. We obtained ∼147 ng/uL of RNA with a RIN greater than 7 for all samples.

### RNA alignment and gene expression analysis

After the quality check using fastqc, data were trimmed using TrimGalore keeping reads with > 30 bp and Phred score > 30. Reads were mapped to the bovine reference genome using STAR and counting using ‘featureCounts’ from Rsubread (version 1.34.7) (Dobin et al., 2015; Liao et al., 2019) with default settings, considering reads with sequencing scores across the total read length that were above 33. According to the ENSEMBL, NCBI and UCSC with NONCODE, RefSeq annotation, mapped reads were summarized at the gene level (Cunningham et al., 2022; Sayers et al., 2022; Zhao et al., 2021). Count summaries were obtained with the ‘featureCounts’ function implemented in Rsubread Package with the strand-specific option and default settings. The final count table for each database had the ENTREZID code aligned for each gene using bitr function from Clusterprofiler package (Yu et al., 2012). After that, using ENSEMBL count table as reference, all repeated ENTREZID in NCBI count table were removed and finally the two tables were merged. For UCSC + NONCODE, only noncode genes were filtered in this count table. Then, the filtered noncode table was added in the merged ENSEMBL/NCBI table. This table was used to proceed with further analysis.

Reads were filtered using the edgeR default threshold (Robinson et al., 2009), where genes with more than three reading counts in at least three samples were used in the DESeq2 model. Differential gene expression analysis was performed using the DESeq2 package (Love et al., 2014). The Benjamini-Hochberg method [“BH”, FDR < 0.05] with an alpha of 0.1 was used to compare the two groups, i.e., BL and CL. PCA (Principal Component Analysis), smear plots (ggplot2), heatmaps (pheatmap), and pathways were performed and constructed using R (Wickham, 2011; Kolde 2022).

